# OCT4 is expressed in extraembryonic endoderm stem (XEN) cell progenitors during somatic cell reprogramming

**DOI:** 10.1101/2024.01.22.576724

**Authors:** Alexandra Moauro, Stephanie L. Hickey, Michael A. Halbisen, Anthony Parenti, Amy Ralston

## Abstract

During development, progenitors of embryonic stem (ES) and extraembryonic endoderm stem (XEN) cells are concomitantly specified within the inner cell mass (ICM) of the mouse blastocyst. Similarly, XEN cells are induced (iXEN cells) alongside induced pluripotent stem (iPS) cells following overexpression of *Oct4*, *Sox2*, *Klf4* and *Myc* (OSKM) during somatic cell reprogramming. It is unclear how or why this cocktail produces both stem cell types, but OCT4 has been associated with non-pluripotent outcomes. In this report, we show that, during OSKM reprogramming, many individual *Oct4-*GFP-expressing cells are fated to become iXEN cells. Interestingly, SKM alone was also sufficient to induce iXEN cell formation, likely via activation of endogenous *Oct4.* These observations indicate that iXEN cell formation is not strictly an artifact of *Oct4* overexpression. Moreover, our results suggest that a pathway to XEN may be an integral feature of establishing pluripotency during reprogramming, as in early embryo development.

## Introduction

During mouse development, progenitors of embryonic stem (ES) and extraembryonic endoderm stem (XEN) cells are co-specified within the inner cell mass (ICM) of the mouse blastocyst. Similarly, somatic cell reprogramming using *Oct4, Sox2, Klf4* and *c-Myc* (OSKM) gives rise to both induced pluripotent stem (iPS) cells (Takahashi and Yamanaka, 2006) and induced XEN (iXEN) cells (Nishimura et al., 2017; Parenti et al., 2016). How and why these four factors produce both iXEN alongside iPS cells is still unclear.

Insight into the roles of reprogramming factors may come from understanding their developmental roles. For instance, *Oct4/Pou5f1* is initially expressed in both pluripotent and extraembryonic endoderm cell types in the mouse blastocyst (Palmieri et al., 1994). Moreover, *Oct4* is required cell-autonomously for both pluripotent epiblast and extraembryonic endoderm development (Frum et al., 2013; Le Bin et al., 2014). These observations are consistent with the observation that OCT4 induces expression of either pluripotency genes or endodermal genes, in a cell type-specific manner (Aksoy et al., 2013).

The dual developmental activities of OCT4 raise the possibility that OCT4 promotes formation of both iPS and iXEN cell types cell-autonomously during reprogramming. Several studies have shown that *Oct4* is expressed in non-pluripotent or partially reprogrammed colonies during reprogramming (Meissner et al., 2007; Mikkelsen et al., 2008; Sridharan et al., 2009). Understanding the identity of cells express *Oct4* during reprogramming is essential to interpretation of any study relying on the use of *Oct4* reporters as readouts of reprogramming outcomes.

Interestingly, OCT4 was recently shown to induce “off-target” reprogramming outcomes, and the exclusion of OCT4 from reprogramming cocktails reportedly improved iPS cell quality (Velychko et al., 2019). These observations raise the possibility that the formation of iXEN cells interferes with optimal formation of iPS cells. Alternatively, formation of iXEN cells could be beneficial or otherwise linked to the formation of iPS cells. Either way, whether iXEN cells arise during SKM reprogramming has not been investigated.

## Results

### Expression of endogenous *Oct4* is observed in presumptive iXEN cell colonies throughout reprogramming

To understand the roles of *Oct4* in somatic cell reprogramming, we used retroviral overexpression of *OSKM* in mouse embryonic fibroblasts (MEFs) (Takahashi and Yamanaka, 2006) carrying *Oct4-eGFP*, a reporter of endogenous *Oct4* expression (Lengner et al., 2007). Consistent with our prior observations (Moauro et al., 2022; Parenti et al., 2016), three main colony morphologies were observed: presumptive iPS cell colonies, which appeared compact and dome-shaped; presumptive iXEN cell colonies, which appeared as patches of mesenchymal cells; and Mixed colonies, which possessed morphological features of both colony types. Interestingly, *Oct4*-eGFP was detected in all three colony subtypes, beginning on day 11 of reprogramming (Fig. 1A). Therefore, endogenous *Oct4* is expressed as cells acquire iXEN cell fate during OSKM reprogramming.

**Figure 1.**
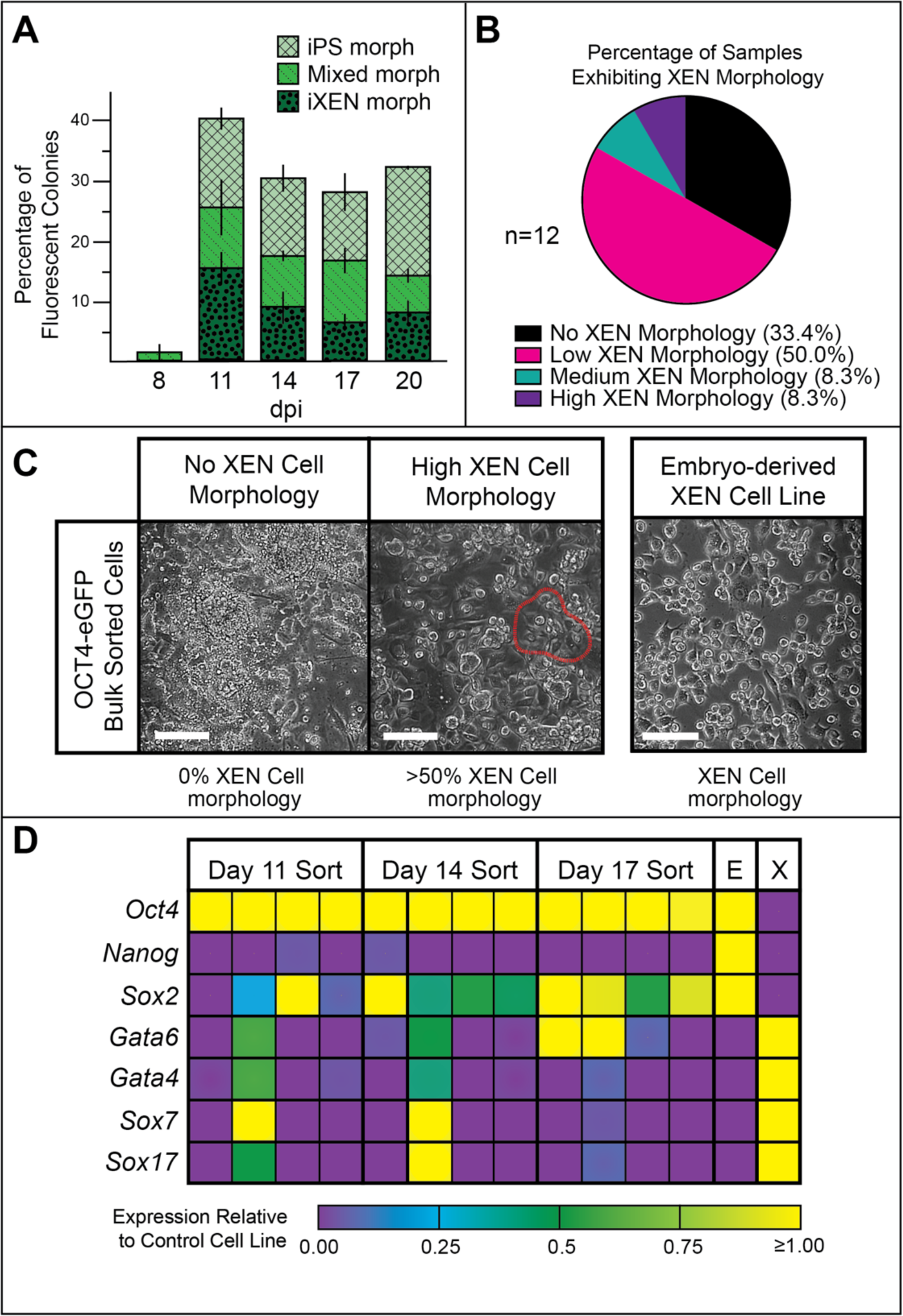
*Oct4* is expressed in presumptive iPS and iXEN cell colonies during OSKM reprogramming. A) Expression of *Oct4*-eGFP was detected within colonies of indicated morphologies during OSKM. Error bars indicate standard error for reprogramming n=3 reprogramming experiments, each with distinct single fetus-derived MEF line, dpi=days post infection. B) Cells expressing *Oct4*-eGFP during reprogramming were FACS-sorted, pooled by time point (n=12 samples), and then morphologically assessed, revealing XEN cell morphology in the majority of samples. C) Examples of morphologies described in panel B (bar=100 µm). D) Gene expression levels in cell pools derived from *Oct4*-eGFP-expressing cells during OSKM reprogramming (n=12), relative to control cell lines (E=ES cells, X=XEN cells) measured by qPCR reveals enrichment of XEN cell markers (*Gata6*, *Gata4*, *Sox7*, and *Sox17*) as well as *Oct4*.

To scrutinize the longer-term, cell-autonomous fates of the *Oct4*-eGFP-expressing cells, we isolated *Oct4*-eGFP-expressing cells on days 11, 14, and 17 of reprogramming by fluorescence-activated cell sorting (n=4 sorts per time point). For our preliminary studies, sorted eGFP-positive cells were pooled, and then allowed to proliferate (n=12 pools). We observed XEN cell-like morphology (Fig. 1B, C) and gene expression (Fig. 1D) among roughly two-thirds of these samples. These observations again suggest that *Oct4* expression is associated with the formation of iXEN cells.

### *Oct4* expression is associated with clonally derived iXEN cell lines

We next evaluated the fates of single *Oct4*-eGFP-expressing cells clonally. We sorted single *Oct4*-eGFP-positive cells into individual wells (n=2,880 sorted cells), and then attempted to expand these as clonal cell lines. From these, we derived 75 cell lines. Among the cell lines that exhibited a singular, stable morphology, around half of these exhibited XEN cell morphology (Fig. 2A). As anticipated, endodermal, but not pluripotency, markers were robustly detected within the clonally derived iXEN cell lies (Fig. 2B, C). Interestingly, OCT4 was initially detected, but was not maintained after prolonged passaging (≥12 passages), suggesting that endogenous OCT4 is expressed transiently during iXEN cell formation. We then evaluated transcriptomes of the *Oct4*-eGFP clonally-derived iXEN cell lines by RNA-sequencing (RNA-seq) (Fig. 2D), which showed that these cell lines were highly similar to both embryo-derived XEN cell lines and iXEN cell lines that had been selected without use of a fluorescent reporter (Moauro and Ralston, 2022; Parenti et al., 2016).

**Figure 2.**
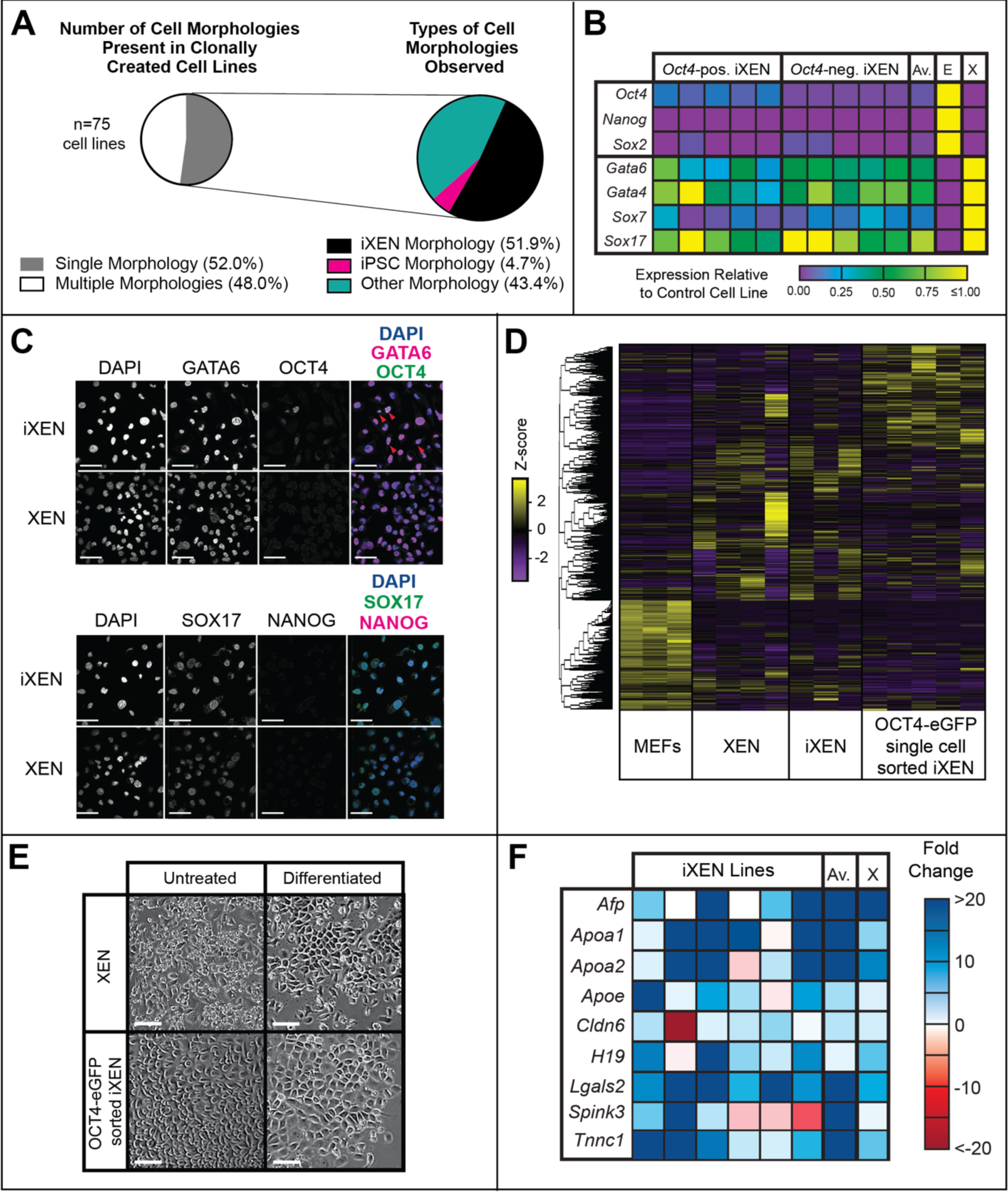
Clonal analysis of cells expressing *Oct4* during OSKM reprogramming reveals iXEN cells progenitor potential. A) Cell lines were clonally established from single *Oct4*-eGFP-positive cells during OSKM reprogramming. The proportion of cells possessing a single morphology was quantified, and the proportion of these that possessed iXEN or iPS cell morphologies is shown. B) Gene expression levels in *Oct4*-eGFP-derived clones (Oct4-pos. iXEN) and our previously characterized iXEN cell lines iXEN cells derived from cell lines not carrying *Oct4-eGFP* (Oct4-neg. iXEN), relative to control cell lines (E=ES cells, X=XEN cells) measured by qPCR reveals individual and average (Av.) enrichment of XEN cell markers (n=5 reprogramming experiments, each with distinct single fetus-derived MEF line). C) Immunofluorescent detection of markers of primitive endoderm and pluripotency shows an exemplary *Oct4*-eGFP-derived clone that expresses low levels of OCT4 not NANOG, bar = 50 µm. D) Heat map showing clustered gene expression in indicated cell lines. Notably, XEN, iXEN, and clonally-derived iXEN cell lines are most highly similar and dissimilar to MEFs. E) Similar morphological changes observed when embryo-derived XEN and clonally-derived iXEN cell lines undergo directed differentiation to visceral endoderm (bar=50µm). F) Markers of visceral endoderm exhibit similar changes in clonally-derived iXEN cell lines, as in XEN cells (X), confirming differentiation of *Oct4-*eGFP-derived iXEN cells to more mature extraembryonic endodermal endpoint (n=6 independent iXEN cell lines).

Finally, we evaluated the developmental potential of the *Oct4*-eGFP clonally-derived iXEN cell lines by directed differentiation to visceral endoderm (Artus et al., 2012; Paca et al., 2012), a more mature derivative of the extraembryonic endoderm. In this assay, iXEN cells underwent the expected morphological and gene expression changes (Fig. 2E, F), consistent with iXEN cell differentiation. Altogether, these observations indicate that OCT4-eGFP-derived iXEN cell lines are highly similar to embryo-derived XEN cell lines, in terms of self-renewal, gene expression, and differentiation. Therefore, expression of endogenous *Oct4* is activated within nascent iXEN cells during OSKM reprogramming.

### *Oct4*-positive cells express primitive endoderm genes during reprogramming

Next, we sought to capture the transcriptional signatures of individual cells undergoing reprogramming, using single-cell RNA-sequencing (scRNA-seq). For this experiment, we focused on 4,570 high quality cell libraries isolated on day 17 of reprogramming. We clustered cell transcriptomes by similarity, yielding eleven stable clusters (Fig. 3A). To characterize these, we compared cluster expression profiles with scRNA-seq expression profiles of mouse early embryonic cell types (Mohammed et al., 2017). This analysis enabled discovery of clusters whose gene expression profile was associated with developmentally relevant cell types (Fig. 3B). We note that Clusters 2 and 5 overlapped strongly and significantly with genes expressed in the primitive endoderm at E4.5 (Fig. 3B, C). This is signification because the primitive endoderm constitutes the XEN cell progenitor population (Kunath et al., 2005). Additionally, Cluster 3 overlapped significantly with visceral endoderm, while Cluster 8 overlapped with E4.5 epiblast and E3.5 ICM genes. As a control, we compared gene expression profiles of ES and XEN cells with the mouse embryo scRNA-seq data, and we observed expected correlations (Fig. 3D). Therefore, both extraembryonic endoderm, as well as pluripotent, signatures are present within a subset of cells undergoing reprogramming.

**Figure 3.**
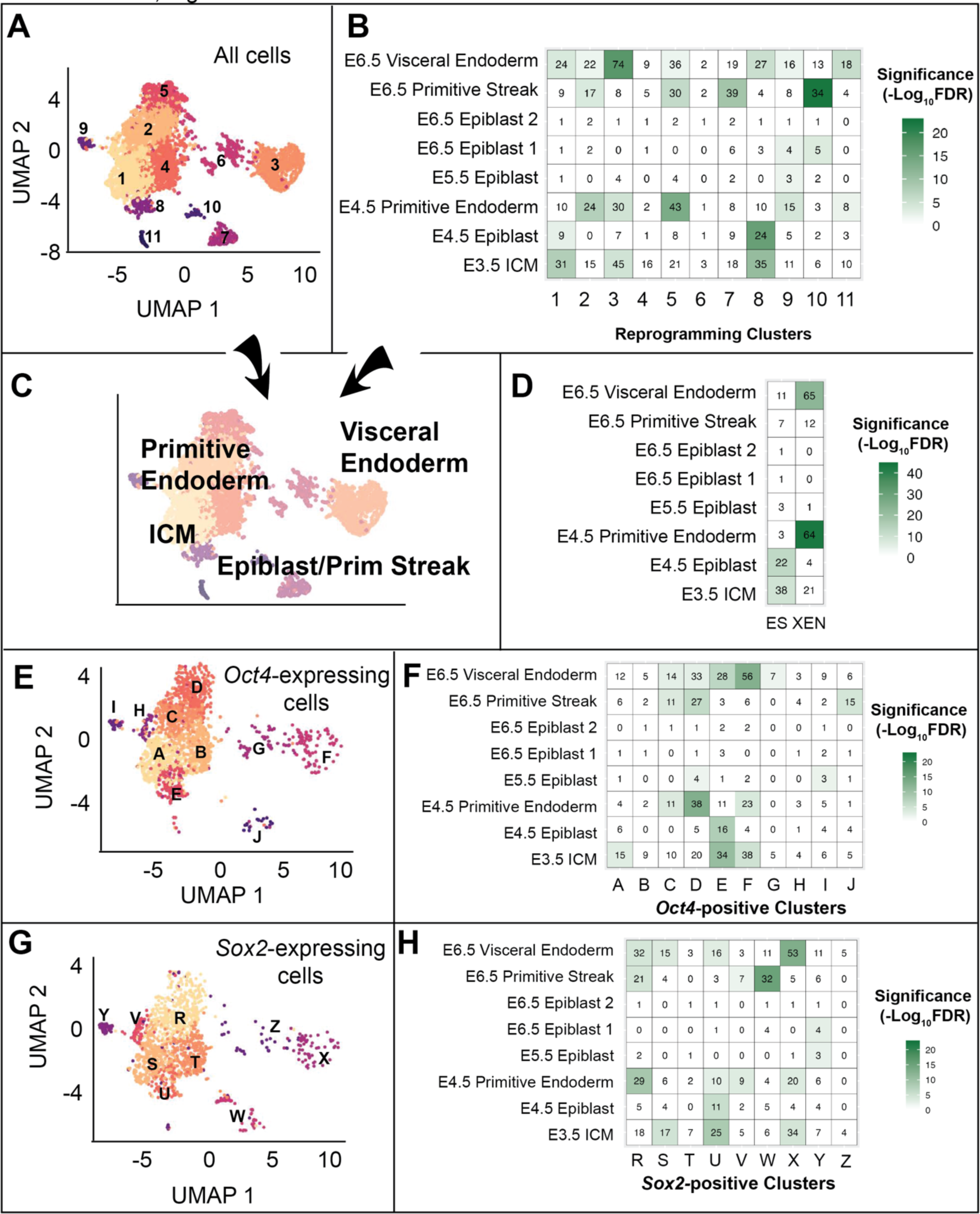
Identification of *Oct4*-expressing cells with primitive endoderm character during OSKM reprogramming by single cell RNA-sequencing. A) UMAP analysis of scRNA-seq data derived from cells on day 17 of OSKM reprogramming cells reveals eleven major clusters (numbered 1-11). B) Overlap of Clusters 1-11 from panel A with single-cell transcriptomes from early embryos (Mohammed et al., 2017) shows significant enrichment (green) of embryonic cell type gene expression within several clusters. ICM = inner cell mass. C) For the purposes of visualization, enrichments from panel B were overlayed on cluster map from panel A. D) Transcriptional profiles from ES and XEN cell bulk RNA-seq were significantly enriched for expected embryonic cell types. E) Cells expressing *Oct4* were reclustered, producing Clusters A-J. F) Overlap of Clusters A-J from panel E with single-cell transcriptomes from early embryos (Mohammed et al., 2017). G) Cells expressing *Sox2* were reclustered, producing Clusters R-Z. F) Overlap of Clusters R-Z from panel G with single-cell transcriptomes from early embryos (Mohammed et al., 2017).

Next, we focused on the individual cells in which *Oct4* expression could be detected by scRNA-seq. Reclustering the *Oct4*-positive cell transcriptomes produced ten major clusters (Fig. 3E). We noted clusters with significant similarity to ICM, E4.5 epiblast, and E4.5 primitive endoderm (Fig. 3F). As a comparison, we reclustered *Sox2*-positive cell transcriptomes (Fig. 3G), which produced similar results (Fig. 3H). These observations are consistent with the developmental roles of OCT4 and SOX2 in promoting both pluripotent and extraembryonic endoderm cell types in the blastocyst (Frum et al., 2013; Le Bin et al., 2014; Wicklow et al., 2014), and reinforce the notion that cells with extraembryonic endoderm transcriptional signature express *Oct4* and exist among cells undergoing reprogramming.

### SKM induce formation of iXEN cells

A recent report demonstrated that *Oct4* can be omitted from the reprogramming cocktail to produce mouse iPS cells, as long as SKM are delivered by a dox-inducible lentivirus, rather than the traditional, retrovirus approach (Velychko et al., 2019). Lentiviral delivery of SKM, while slower and less efficient, yielded a more specific population of higher quality iPS cells, which led to the conclusion that OCT4 is responsible for undesired reprogramming outcomes. These observations raised the possibility that iXEN cell formation is an artifact of including Oct4 in the reprogramming cocktail.

To investigate this possibility, we used the dox-inducible lentiviral system to overexpress either OSKM or SKM in MEFs. We observed dramatically reduced colony number in SKM, compared with OSKM reprogramming (Fig. 4A), consistent with prior observations (Velychko et al., 2019). Interestingly, we observed both iPS and iXEN cell colonies during lentiviral SKM reprogramming, based on morphological criteria (Fig. 4B). Notably, the yield of putative iPS cell colonies was increased relative to the yield of iXEN cell colonies in the absence of exogenous *Oct4*. As we have previously observed, the ratio of iXEN to iPS cell colonies was 3:1 by day 20 of retroviral OSKM reprogramming (Parenti et al., 2016). By contrast, the ratio of iXEN to iPS cell colonies was 1:1 by day 20 of lentiviral SKM reprogramming. These observations suggest that while SKM favors formation of iPS cells, iXEN cells are still produced using this reprogramming cocktail.

**Figure 4.**
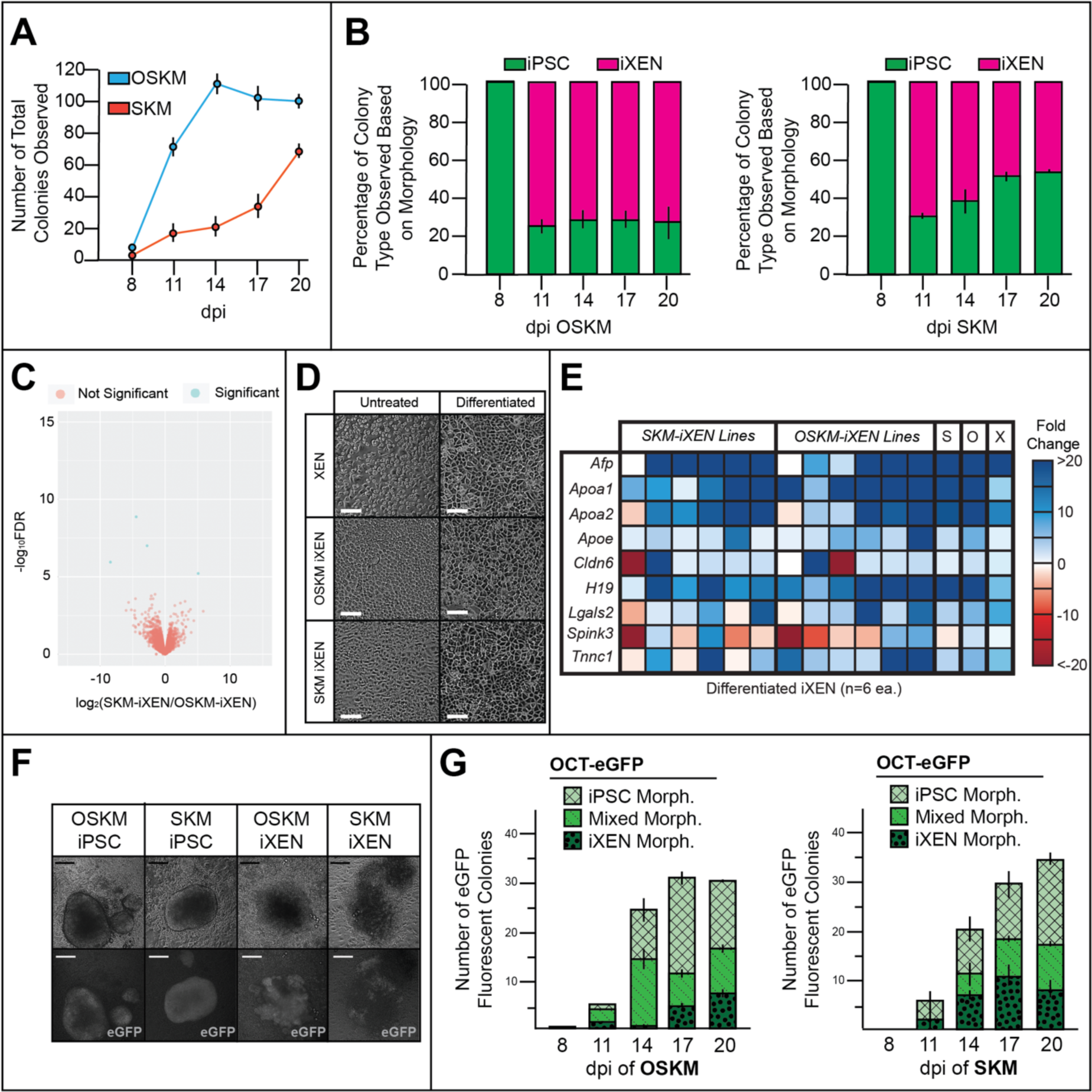
Formation of iXEN cells in the absence of exogenous *Oct4*. A) Quantification of total colonies observed during lentiviral reprogramming of MEFs using OSKM or SKM (dpi=days post infection). B) Relative yield of colonies possessing iPS or iXEN cell morphology during lentiviral OSKM and SKM reprogramming (dpi=days post infection, n=3 reprogramming experiments, each with distinct single fetus-derived MEF line). C) Volcano plot comparing differentially expressed genes between OSKM and SKM derived iXEN lines show that lines are undistinguishable with only four genes differentially expressed (n=5). D) Similar morphological changes observed when embryo-derived XEN, OSKM-derived, and SKM-derived iXEN cell lines underwent directed differentiation to visceral endoderm (bar =50 µm). E) Markers of visceral endoderm markers exhibit anticipated expression changes in OSKM-derived and SKM-derived iXEN cell lines, confirming differentiation to more mature extraembryonic endodermal endpoint (S=Average for SKM-induced iXEN cell lines, O=average for OSKM-induced iXEN cell lines, X=XEN cells, n=6 reprogramming experiments, each with distinct single fetus-derived MEF line). F) Images of *Oct4*-eGFP expression within colonies bearing iPS and iXEN cell morphologies lentivirally reprogrammed using OSKM or SKM (bar=100µm). G) Quantification of *Oct4-*eGFP expression within colony subtypes during lentiviral reprogramming using OSKM or SKM (n=3 reprogramming experiments each, each with distinct single fetus-derived MEF line).

To confirm the identity of the presumptive iXEN cell colonies formed during SKM reprogramming, we established stable stem cell lines. We observed the appropriate and homogenous morphologies and expression of ES and XEN cell markers in among these cell lines (Supplemental Fig. 4). Using RNA-seq, significant differences in gene expression were detected for only four genes (Fig. 4C), indicating that iXEN cells derived using OSKM or SKM are highly transcriptionally similar. These four genes (*Psca*, *Clic6*, *Rpl21*, and *Rps26-ps1*) have no known roles in XEN cells. Therefore, SKM-induced iXEN cell lines are highly similar to OSKM-induced iXEN and embryo-derived XEN cell lines in terms of morphology and gene expression.

Finally, we evaluated the stem cell properties of the SKM and OSKM-induced iXEN cell lines. Both OSKM and SKM-induced iXEN cell lines were capable of continuous expansion (≥20 passages), consistent with self-renewal. Additionally, we observed that SKM and OSKM-induced iXEN cell lines exhibited expected changes in morphology and gene expression during directed differentiation to visceral endoderm-like cells (Fig. 4D, E). We conclude that iXEN cells derived in either the presence or absence of exogenous OCT4 are highly comparable to embryo-derived XEN cells.

### SKM induce expression of endogenous *Oct4* in iXEN cell progenitors

SKM has been reported to induce expression of endogenous *Oct4* in iPS cells (Velychko et al., 2019), but its association with iXEN cells is unknown. We reprogrammed MEFs carrying *Oct4-eGFP* using lentiviral reprogramming. Indeed, we observed expression of *Oct4*-eGFP in presumptive iXEN cell colonies during OSKM, as well as SKM, lentiviral reprogramming (Fig. 4F, G). We conclude that endogenous *Oct4* expression is associated with formation of iXEN cells in all conditions tested here.

## Discussion

We have shown that *Oct4* expression is associated with multiple types of stem cell during mouse cell reprogramming. This is significant because *Oct4* is still a commonly used readout of pluripotency and efficiency of iPS cell formation during reprogramming, yet the presence of fluorescent iXEN cells could interfere with accurate quantification of iPS cell efficiency, depending on the reprogramming protocol used.

We note that OCT4 was previously proposed to exhibit activity that is “off-target” in the context of mouse cell reprogramming (Velychko et al., 2019). However, it is unclear whether off-target effects represent gene expression noise or a distinct, biologically relevant transcriptional role for OCT4. Arguing for the latter, OCT4 promotes primitive endoderm gene expression cell-autonomously in the blastocyst (Frum et al., 2013; Le Bin et al., 2014). In addition, OCT4 binding patterns differ between pluripotent and extraembryonic endoderm cells, promoting either pluripotency or endodermal gene expression (Aksoy et al., 2013).

An intriguing yet unanswered question is how OCT4 achieves its two roles, whether ectopically or endogenously expressed. Our study raises several prospects. First, MEF subtypes could exist with differential enrichment of cofactors that impact OCT4 activity. Consistent with this, SOX2 is thought to lead OCT4 to pluripotency gene expression targets, while SOX17 leads OCT4 to endodermal targets in cell lines (Aksoy et al., 2013). A second and related possibility to explain how OCT4 might accomplish its dual roles is that MEF subtypes could possess differing chromatin landscapes, enabling differential access (or pioneering potential) to OCT4 near endodermal versus pluripotency gene regulatory regions. If true for MEFs, the same could be true for adult fibroblasts, since iXEN cells can be derived from adult fibroblasts with similar efficiency (Parenti et al., 2016). However, very little is known about chromatin state during the divergence of pluripotent and XEN cell states during either embryogenesis or somatic cell reprogramming.

A third way in which OCT4 could induce the two types of stem cell during reprogramming might involve regionalized intercellular interactions that randomly establish local differences in signaling pathways and/or OCT4 activity. Given our observation that OCT4 is expressed in an ICM-like cluster of cells resolved by since cell RNA-seq, the possibility of local intercellular communication is particularly intriguingly. This is because in the ICM, OCT4 and SOX2 drive expression of FGF4 in pluripotent cells, which induces primitive endoderm gene expression non cell-autonomously (Frum et al., 2013; Kang et al., 2013; Krawchuk et al., 2013; Wicklow et al., 2014; Yamanaka et al., 2010; Yuan et al., 1995).

Finally, we note that resolving how many distinct cell types exist during reprogramming continues to present a challenge to the field. This is important for several reasons, among which is our capacity to understand the origins of iXEN cells during reprogramming. We previously used lineage tracing to show that iXEN cells are not derived from iPS cells, nor vice-versa, during OSKM reprogramming (Parenti et al., 2016). However, we do not yet know whether iPS and iXEN cells derive directly from MEFs, or whether some or all MEFs pass through an intermediate ICM-like state capable of giving rise to either iPS or iXEN cells. If true, this could indicate that MEF reprogramming is even more like mouse embryogenesis than previously recognized.

## Experimental Procedures

### Mouse Strains

All animal work conformed to the guidelines and regulatory standards of Michigan State University Institutional Animal Care and Use Committee. The following alleles were The *Pou5f1^tm2Jae^*allele (Lengner et al., 2007) was maintained as a heterozygote in a CD-1 background.

### Mouse Embryonic Fibroblast (MEF) Preparation and Reprogramming

MEF lines were established from individual E13.5 embryos of differing genotypes, and individual lines were reprogrammed as described (Moauro and Ralston, 2022; Parenti et al., 2016). See Supplemental Methods for detailed description.

### Fluorescence Activated Cell Sorting

Cells were harvested, passed through a 40 mm filter, and then sorted by 488 nm laser and BD Influx. Gating was facilitated by using ES cells carrying *Oct4-eGFP*, and cells that were eGFP-positive and DAPI-negative were sorted onto irradiated MEFs in 12-well plates (for pooled cell studies) or 96-well plates (for single cell studies).

### Immunofluorescence and Confocal Microscopy

Cells were plated the day before staining on confocal grade plastic (ibidi) and fixed with 4% formaldehyde (Polysciences) for 10 min, permeabilized with 0.5% Triton X-100 (Millipore Sigma), then blocked in 10% fetal bovine serum (Hyclone) with 0.1% Triton X-100 at 4°C then incubated overnight in primary antibody at 4°C. Antibodies are listed in Supplemental Experimental Procedures. Imaging was performed using an Olympus FluoView FV1000 Confocal Laser Scanning Microscope system with a 40x UPLFL oil lens 1.30 NA or 60x UPLFL oil lens 1.25 NA.

### Directed XEN cell Differentiation

Directed XEN cell differentiation to visceral endoderm was achieved as previously described (Artus et al., 2012; Paca et al., 2012; Parenti et al., 2016). Wells were pre-treated with Poly-L-ornithine (Sigma) for 30 min at room temperature, and then with 0.15 μg/cm^2^ Laminin (Sigma). Cells were plated at a density of ∼11,000 cells/cm^2^ in N2B27 Medium [50% DMEM-F12 (Invitrogen) + 50% Neural Basal Medium (Invitrogen) + N2 Medium (Invitrogen, 100x) + B27 (Invitrogen, 50x) + Pen/Strep (10,000 units each), beta-mercaptoethanol (55 mM)], and were then cultured overnight at 37°C and 5% CO2. On days 2, 4, 6 and 8 the culture medium was replaced with fresh N2B27 + 50 ng/μL BMP4 (R&D Systems) and analyses performed on day 9.

### Transcriptional Analyses and Data Availability

For procedures, please see Supplemental Experimental Procedures. The data discussed in this publication have been deposited in NCBI’s Gene Expression Omnibus (Edgar et al., 2002) and are accessible through GEO Series accession number GSE244818 at https://www.ncbi.nlm.nih.gov/geo/query/acc.cgi and on Zenodo at DOI: DOI: https://doi.org/10.5281/zenodo.8388276.

## Acknowledgements

This study was supported by Award R35 GM131759 to AR and the BMB TEAM UP Award to SLH. We would like to acknowledge Dr. R. William Henry, the Michigan State University Flow Cytometry Core, and the Michigan State University Research Technology and Support Facility Genomics Core for technical support.

## Author Contributions

Conceptualization was provided by AR, with input from AM. Investigations were pursued by AM and AP. Formal analysis was performed by AM, SLH, and MAH. Data curation was performed by SLH and MAH. Writing – original draft was prepared by AM and SLP – review & editing by AM, SLP, and AR. Visualization was prepared by AM and AR. Supervision, project administration and funding acquisition was performed by AR.

---

**Supplemental to Fig. 4.**
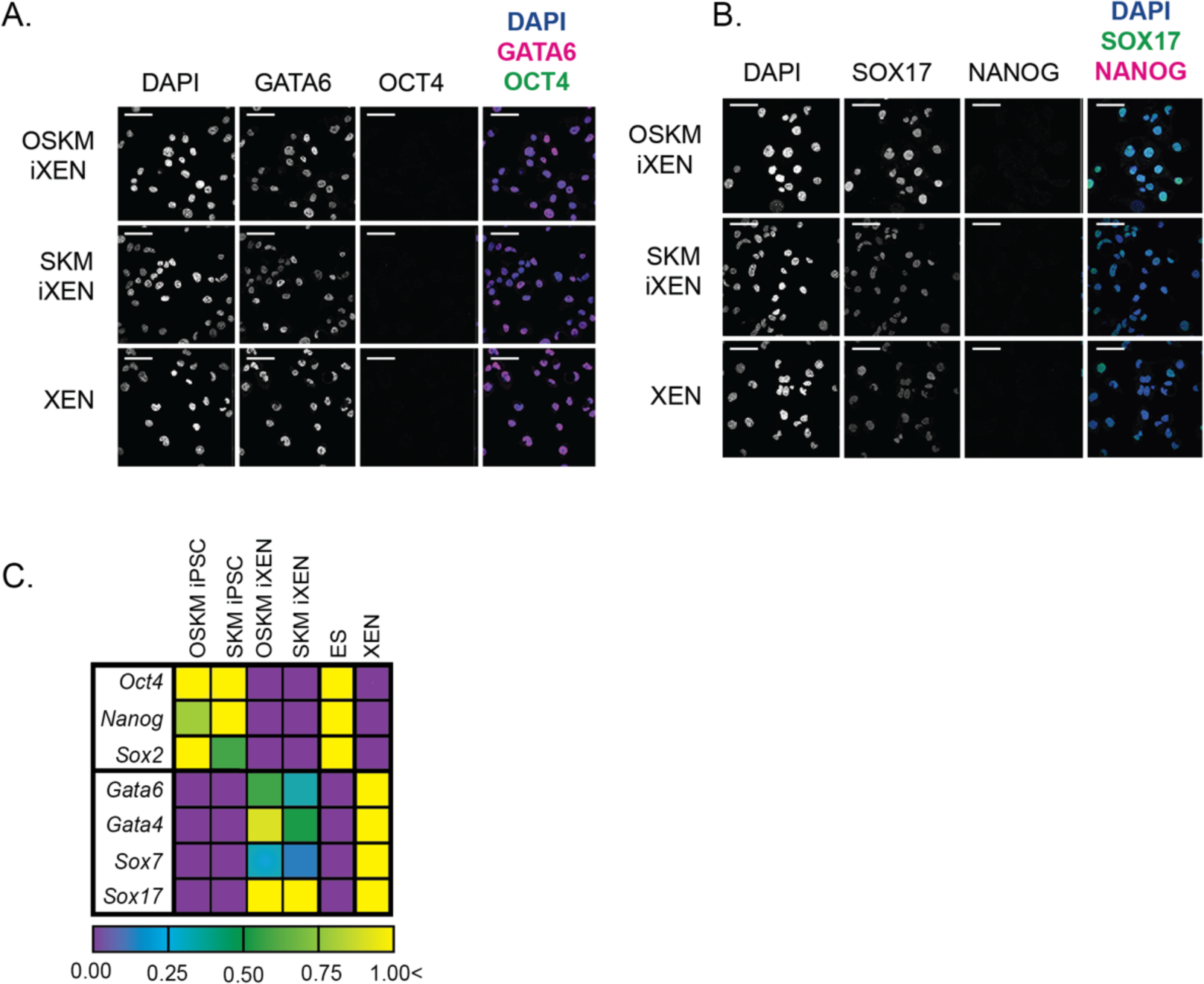
SKM-derived iXEN cells express XEN cell markers. A-B) Immunofluoresence localization of indicated markers in SKM and OSKM lentivirally-induced iXEN cell lines (bar=50 µm). C) Average relative levels of endodermal and pluripotency markers in SKM and OSKM iXEN cell lines by qPCR (n>3 reprogramming experiments, each with distinct single fetus-derived MEF line per category).

## References

Aksoy, I., Jauch, R., Chen, J., Dyla, M., Divakar, U., Bogu, G.K., Teo, R., Leng Ng, C.K., Herath, W., Lili, S., et al. (2013). Oct4 switches partnering from Sox2 to Sox17 to reinterpret the enhancer code and specify endoderm. EMBO J 32, 938–953. 10.1038/emboj.2013.31.

Artus, J., Douvaras, P., Piliszek, A., Isern, J., Baron, M.H., and Hadjantonakis, A.K. (2012). BMP4 signaling directs primitive endoderm-derived XEN cells to an extraembryonic visceral endoderm identity. Developmental Biology 361, 245–262. 10.1016/j.ydbio.2011.10.015.

Edgar, R., Domrachev, M., and Lash, A.E. (2002). Gene Expression Omnibus: NCBI gene expression and hybridization array data repository. Nucleic Acids Res 30, 207–210. 10.1093/nar/30.1.207.

Frum, T., Halbisen, M.A., Wang, C., Amiri, H., Robson, P., and Ralston, A. (2013). Oct4 Cell-autonomously promotes primitive endoderm development in the mouse blastocyst. Developmental Cell 25, 610–622. 10.1016/j.devcel.2013.05.004.

Kang, M., Piliszek, A., Artus, J., and Hadjantonakis, A.K. (2013). FGF4 is required for lineage restriction and salt-and-pepper distribution of primitive endoderm factors but not their initial expression in the mouse. Development 140, 267–279. 10.1242/dev.084996.

Krawchuk, D., Honma-Yamanaka, N., Anani, S., and Yamanaka, Y. (2013). FGF4 is a limiting factor controlling the proportions of primitive endoderm and epiblast in the ICM of the mouse blastocyst. Dev Biol 384, 65–71. 10.1016/j.ydbio.2013.09.023.

Kunath, T., Arnaud, D., Uy, G.D., Okamoto, I., Chureau, C., Yamanaka, Y., Heard, E., Gardner, R.L., Avner, P., and Rossant, J. (2005). Imprinted X-inactivation in extra-embryonic endoderm cell lines from mouse blastocysts. Development 132, 1649–1661. 10.1242/dev.01715.

Le Bin, G.C., Muñoz-Descalzo, S., Kurowski, A., Leitch, H., Lou, X., Mansfield, W., Etienne-Dumeau, C., Grabole, N., Mulas, C., Niwa, H., et al. (2014). Oct4 is required for lineage priming in the developing inner cell mass of the mouse blastocyst. Development 141, 1001–1010. 10.1242/dev.096875.

Lengner, C.J., Camargo, F.D., Hochedlinger, K., Welstead, G.G., Zaidi, S., Gokhale, S., Scholer, H.R., Tomilin, A., and Jaenisch, R. (2007). Oct4 Expression Is Not Required for Mouse Somatic Stem Cell Self-Renewal. Cell Stem Cell 1, 403–415. 10.1016/j.stem.2007.07.020.

Meissner, A., Wernig, M., and Jaenisch, R. (2007). Direct reprogramming of genetically unmodified fibroblasts into pluripotent stem cells. Nat Biotechnol 25, 1177–1181. nbt1335 [pii] 10.1038/nbt1335.

Mikkelsen, T.S., Hanna, J., Zhang, X., Ku, M., Wernig, M., Schorderet, P., Bernstein, B.E., Jaenisch, R., Lander, E.S., and Meissner, A. (2008). Dissecting direct reprogramming through integrative genomic analysis. Nature 454, 49–55. 10.1038/nature07056.

Moauro, A., and Ralston, A. (2022). Distinguishing Between Endodermal and Pluripotent Stem Cell Lines During Somatic Cell Reprogramming. Methods Mol Biol 2429, 41–55. 10.1007/978-1-0716-1979-7_4.

Moauro, A., Kruger, R.E., O’Hagan, D., and Ralston, A. (2022). Fluorescent Reporters Distinguish Stem Cell Colony Subtypes During Somatic Cell Reprogramming. Cell Reprogram 24, 353–362. 10.1089/cell.2022.0071.

Mohammed, H., Hernando-Herraez, I., Savino, A., Scialdone, A., Macaulay, I., Mulas, C., Chandra, T., Voet, T., Dean, W., Nichols, J., et al. (2017). Single-Cell Landscape of Transcriptional Heterogeneity and Cell Fate Decisions during Mouse Early Gastrulation. Cell Reports 20, 1215–1228. 10.1016/j.celrep.2017.07.009.

Nishimura, T., Unezaki, N., Kanegi, R., Wijesekera, D.P.H., Hatoya, S., Sugiura, K., Kawate, N., Tamada, H., Imai, H., and Inaba, T. (2017). Generation of Canine Induced Extraembryonic Endoderm-Like Cell Line That Forms Both Extraembryonic and Embryonic Endoderm Derivatives. Stem Cells and Development 26, 1111–1120. 10.1089/scd.2017.0026.

Paca, A., Séguin, C.A., Clements, M., Ryczko, M., Rossant, J., Rodriguez, T.A., and Kunath, T. (2012). BMP signaling induces visceral endoderm differentiation of XEN cells and parietal endoderm. Developmental Biology 361, 90–102. 10.1016/j.ydbio.2011.10.013.

Palmieri, S.L., Peter, W., Hess, H., and Schöler, H.R. (1994). Oct-4 Transcription Factor Is Differentially Expressed in the Mouse Embryo during Establishment of the First Two Extraembryonic Cell Lineages Involved in Implantation. Developmental Biology 166, 259–267. 10.1006/dbio.1994.1312.

Parenti, A., Halbisen, M.A., Wang, K., Latham, K., and Ralston, A. (2016). OSKM Induce Extraembryonic Endoderm Stem Cells in Parallel to Induced Pluripotent Stem Cells. Stem Cell Reports 6, 447–455. 10.1016/j.stemcr.2016.02.003.

Sridharan, R., Tchieu, J., Mason, M.J., Yachechko, R., Kuoy, E., Horvath, S., Zhou, Q., and Plath, K. (2009). Role of the murine reprogramming factors in the induction of pluripotency. Cell 136, 364–377. 10.1016/j.cell.2009.01.001.

Takahashi, K., and Yamanaka, S. (2006). Induction of Pluripotent Stem Cells from Mouse Embryonic and Adult Fibroblast Cultures by Defined Factors. Cell 10.1016/j.cell.2006.07.024.

Velychko, S., Adachi, K., Kim, K., Hou, Y., MacCarthy, C.M., Wu, G., and Schöler, H.R. (2019). Excluding Oct4 from Yamanaka Cocktail Unleashes the Developmental Potential of iPSCs. Cell Stem Cell 1–17. 10.1016/j.stem.2019.10.002.

Wicklow, E., Blij, S., Frum, T., Hirate, Y., Lang, R.A., Sasaki, H., and Ralston, A. (2014). HIPPO Pathway Members Restrict SOX2 to the Inner Cell Mass Where It Promotes ICM Fates in the Mouse Blastocyst. PLoS Genetics 10, e1004618. 10.1371/journal.pgen.1004618.

Yamanaka, Y., Lanner, F., and Rossant, J. (2010). FGF signal-dependent segregation of primitive endoderm and epiblast in the mouse blastocyst. Development 137, 715–724. 137/5/715 [pii] 10.1242/dev.043471.

Yuan, H., Corbi, N., Basilico, C., and Dailey, L. (1995). Developmental-specific activity of the FGF-4 enhancer requires the synergistic action of Sox2 and Oct-3. Genes Dev. 9, 2635–2645. 10.1101/gad.9.21.2635.

