## Supplemental Methods for "OCT4 is expressed in extraembryonic endoderm stem (XEN) cell progenitors during somatic cell reprogramming"

**Supplemental Experimental Procedures, Moauro et al.**

**MEFs and Reprogramming**

MMLV-derived retrovirus and lentivirus were produced by transfecting 293T cells with pCL-ECO and pMXs plasmids or pMD2.g, psPAX2 and FUW plasmids, respectively. pMXs plasmids contained either *Oct4*, *Klf4*, *Sox2* or *cMyc* cDNAs (Addgene). FUW plasmids contained either *rtta*, *Oct4*, *Klf4*, *Sox2* or *cMyc* housed under the control of a tetracycline on promoter (Addgene). Transfected 293T cell supernatant was harvested 48 hours later. mCherry virus was made in conjunction with all viral preps and used to infect CD-1 MEFs to determine viral titers. Viral preps were stored at -80 ºC. For retroviral and lentiviral reprogramming (Takahashi and Yamanaka, 2006) MEFs are plated the day before at a density of 50-100 cells/$mm^2$. Virus was then added at a multiplicity of infection (MOI) of 1 with polybrene and incubated for 24 hrs. The following day, medium was replaced with MEF medium, followed by Reprogramming Medium 1 [DMEM (Invitrogen), 0.1 mM Beta-mercaptoethanol, 2 mM Glutamax, 1X Non-essential amino acids, 100 U/mL Penicillin/streptomycin, 15% Fetal bovine serum (FBS; Hyclone), 10 ng/mL Leukemia Inhibitory Factor (LIF)(Millipore Sigma)] on days 2 and 4. Medium was then replaced with Reprogramming Medium 2 [DMEM (Invitrogen), 0.1 mM Beta-mercaptoethanol, 2 mM Glutamax, 1X Non-essential amino acids, 100 U/mL Penicillin/streptomycin, 15% Knockout Serum Replacement (KOSR; Invitrogen), 10 ng/mL Leukemia Inhibitory Factor (LIF)] on day 6 and every other day there after until the end of the experiment. For lentiviral reprogramming, reprogramming medium was supplemented with 2 mg/mL of doxycycline. Cells were grown at 37°C and 5% $CO_2$. At the end of reprogramming, cells were manually picked and expanded ≥12 passages to create stable cell lines.

**Antibodies used for immunofluorescence**

Primary antibodies used were SOX17 (R&D, AF1924, 1:800 dilution), NANOG (Reprocell, RCAB002P_F, 1:400 dilution), OCT4 (Santa Cruz, sc-5279, 1:100 dilution) and GATA6 (R&D,

AF1700, 1:100 dilution). The next day, cells were washed for 30 min in block, then stained for 1 hr with donkey-anti-rabbit Alexa647 (Jackson ImmunoResearch, 711-606-152, 1:400 dilution), Bovine-anti-goat DyLight488 (Jackson ImmunoResearch, 805-005-180, 1:400 dilution), and donkey-anti-mouse DyLight649 (Jackson ImmunoResearch, 715-495-150, 1:400 dilution) secondary antibody. Following staining, cells were again washed for 30 min in block, then stained for 5 min in DAPI (Sigma, D9542-1MG; 1:1000 dilution).

**Transcriptional analysis**

RNA was harvested using 1:6 chloroform to Trizol (Invitrogen), and then1 μg RNA was reverse transcribed to create cDNA using QuantiTect Reverse Transcription Kit (Qiagen), following manufacturer instructions. For qPCR, cDNA levels were measured in quadruplicate, relative to ES or XEN cell levels using a Lightcycler 480 (Roche), according to manufacturer guidelines. For RNA-seq, cell lines were cultured for at least three passages in XEN cell media [70% feeder conditioned media (RPMI (Invitrogen) + 20% FBS (Hyclone) + 100 μM beta-mercaptoethanol + 2 mM glutamax + 1mM sodium pyruvate + 50 μg/mL penicillin/streptomycin) supplemented with 0.025 ng/mL FGF4 (R&D Systems) and 0.001 U/mL Heparin (Sigma-Aldrich)]. Libraries were prepared from 1 μg of RNA using Illumina Stranded mRNA Library Preparation kit, and libraries were sequenced using an Illumina NovaSeq 6000, to a depth of 50-90 million with 50 bp pair-end end reads per sample.

Adapter sequences were removed with Trimmomatic/0.32 (Bolger et al., 2014), and then trimmed raw sequencing reads were aligned to mm10 (https://genome.ucsc.edu/) with hisat2/2.1.0 (Kim et al., 2015, 2019; Pertea et al., 2016; Zhang et al., 2021), and were then counted with HTSeq/0.11.2-Python-3.6.6 (Putri et al. 2021, https://www.python.org/). Experimental design parameters, including sample size and sequencing depth were based on prior analysis (Ching et al., 2014). Sequence quality was evaluated before and after read mapping with FastQC/0.11.7-Java-1.8.0_162 (Wingett et al. 2018) and mapping rates ranged from 85%-99%. Transcripts with low abundance (without ≥10 counts per million in at least 3 XEN, 3 Parenti iXEN, 5 SKM iXEN, 5 OSKM iXEN and 5 OCT4-eGFP iXEN samples) were removed, differential gene expression analysis was completed with EdgeR 3.24.3 (Chen et al., 2016; McCarthy et al., 2012; Robinson et al., 2009). Volcano plots were generated using the ggplot2 3.3.0 package (Wickham, 2016). Pairwise Spearman correlations (Glasser and Winter, 1961, (Spearman, 2010) were calculated for each sample, and the heatmap.2 function of ggplots 3.0.1 (Warnes et al., 2016) were used to generate heat maps. All subsequent bioinformatic analyses were performed in R/3.3.1 (R Core Team 2018). Raw and processed RNA sequencing files used in this study will be archived and available from the Gene Expression Omnibus database.

For single cell RNA-seq, samples were passaged one day before submission to remove dead cells and cell debris. On the day of analysis, cells were harvested and filtered as described above. Submitted samples contained <1% of cell clumps and a cell viability of >95%. Paired-end libraries were prepared using the 10x Genomics Single Cell 3' V3.1 kit and sequenced on an Illumina NovaSeq 6000. Base calling was performed using Illumina Real Time Analysis (RTA), and the output of RTA was demultiplexed and converted to FastQ format with Illumina Bcl2fastq v2.20.0. After demultiplexing and FastQ conversion, cellrangercount v6.1.1 was used for alignment to the mm10 reference transcriptome (included with cellranger), cell detection and UMI counting.

Cells with <10% of reads coming from mitochondrial genes, >5,000 UMIs, and >1000 detected genes were used in the analysis, for a total of 4,507 cells. Transcriptional analysis was completed using R v4.1.0 (R Core Team, 2021) with tools from Seurat v4.1.0 (Hao et al., 2021). UMI counts were normalized using SCTransform, regressing the percent of reads coming from mitochondrial genes. To define cell clusters, we used the FindNeigbors function with the first 30 principal components followed by the FindClusters function with a resolution of 0.70. The clusters were visualized using UMAP performed on the first 30 principal components.

We identified 1,437 *Oct4*-positive and 931 Sox2-positive cells among 4,570 total. For each subset, UMI counts were re-normalized and cells were re-clustered as described above. In all analyses, cluster enriched genes were identified using the FindAllMarkers function. Cluster-enriched genes included those with a log2 fold change threshold >0.25 expressed in at least 10% of the cells in either the cluster of interest with adjusted p<0.001. Enriched genes were compared with cluster-enriched genes from mouse early embryos (Mohammed et al., 2017) and genes differentialy expressed in ES and XEN cell bulk RNA-seq using a hypergeometric test. P-values were corrected for multiple comparisons using the Benjamini-Hochberg procedure. ES and XEN differentially expressed genes were identified using Illumina MouseWG-6_V2 expression BeadChip array data generated by Wamaitha *et. al.* (Wamaitha et al., 2015). The non-normalized data was downloaded from GEO (GSE69321), and data derived from mESC and eXEN cells were extracted and processed using the R/Bioconductor package, beadarray (Dunning et al., 2007). Briefly, probes with quality described as "No Match" or "Bad" in the illuminaMousev2 annotation database were removed from the analysis, and for each gene assayed with multiple probes, the probe with the highest average expression across samples was selected (Miller et al., 2011). The expression data was quantile normalized and $\log_2$ transformed. Differential expression analysis between ES and XEN cells was performed using the limma R/Bioconductor package (Ritchie et al., 2015). Cluster-enriched genes from mouse early embryos (Mohammed et al., 2017) were compared with genes differentially expressed (adjusted p-value < .01) in

ES and XEN cells using a hypergeometric test. P-values were corrected for multiple comparisons using the Benjamini-Hochberg procedure.

**References**


Bolger, A.M., Lohse, M., and Usadel, B. (2014). Trimmomatic: A flexible trimmer for Illumina sequence data. Bioinformatics *30*, 2114–2120. https://doi.org/10.1093/bioinformatics/btu170.

Chen, Y., Lun, A.T.L., and Smyth, G.K. (2016). From reads to genes to pathways: differential expression analysis of RNA-Seq experiments using Rsubread and the edgeR quasi-likelihood pipeline. F1000Research *5*, 1438. https://doi.org/10.12688/f1000research.8987.1.

Ching, T., Huang, S., and Garmire, L.X. (2014). Power analysis and sample size estimation for RNA-Seq differential expression. Rna *20*, 1684–1696. https://doi.org/10.1261/rna.046011.114.

Dunning, M.J., Smith, M.L., Ritchie, M.E., and Tavaré, S. (2007). beadarray: R classes and methods for Illumina bead-based data. Bioinformatics *23*, 2183–2184. https://doi.org/10.1093/bioinformatics/btm311.

Hao, Y., Hao, S., Andersen-Nissen, E., Mauck, W.M., Zheng, S., Butler, A., Lee, M.J., Wilk, A.J., Darby, C., Zager, M., et al. (2021). Integrated analysis of multimodal single-cell data. Cell *184*, 3573-3587.e29. https://doi.org/10.1016/j.cell.2021.04.048.

Kim, D., Langmead, B., and Salzberg, S.L. (2015). HISAT: a fast spliced aligner with low memory requirements. Nature Methods *12*, 357–360. https://doi.org/10.1038/nmeth.3317.

Kim, D., Paggi, J.M., Park, C., Bennett, C., and Salzberg, S.L. (2019). Graph-based genome alignment and genotyping with HISAT2 and HISAT-genotype. Nature Biotechnology *37*, 907–915. https://doi.org/10.1038/s41587-019-0201-4.

McCarthy, D.J., Chen, Y., and Smyth, G.K. (2012). Differential expression analysis of multifactor RNA-Seq experiments with respect to biological variation. Nucleic Acids Research *40*, 4288–4297. https://doi.org/10.1093/nar/gks042.

Miller, J.A., Cai, C., Langfelder, P., Geschwind, D.H., Kurian, S.M., Salomon, D.R., and Horvath, S. (2011). Strategies for aggregating gene expression data: The collapseRows R function. BMC Bioinformatics *12*, 322. https://doi.org/10.1186/1471-2105-12-322.

Mohammed, H., Hernando-Herraez, I., Savino, A., Scialdone, A., Macaulay, I., Mulas, C., Chandra, T., Voet, T., Dean, W., Nichols, J., et al. (2017). Single-Cell Landscape of Transcriptional Heterogeneity and Cell Fate Decisions during Mouse Early Gastrulation. Cell Reports *20*, 1215–1228. https://doi.org/10.1016/j.celrep.2017.07.009.

Pertea, M., Kim, D., Pertea, G.M., Leek, J.T., and Salzberg, S.L. (2016). Transcript-level expression analysis of RNA-seq experiments with HISAT, StringTie and Ballgown. Nature Protocols *11*, 1650–1667. https://doi.org/10.1038/nprot.2016.095.

Ritchie, M.E., Phipson, B., Wu, D., Hu, Y., Law, C.W., Shi, W., and Smyth, G.K. (2015). limma powers differential expression analyses for RNA-sequencing and microarray studies. Nucleic Acids Res *43*, e47. https://doi.org/10.1093/nar/gkv007.

Robinson, M.D., McCarthy, D.J., and Smyth, G.K. (2009). edgeR: A Bioconductor package for differential expression analysis of digital gene expression data. Bioinformatics *26*, 139–140. https://doi.org/10.1093/bioinformatics/btp616.

Spearman, C. (2010). The proof and measurement of association between two things. International Journal of Epidemiology *39*, 1137–1150. https://doi.org/10.1093/ije/dyq191.

Wamaitha, S.E., Del Valle, I., Cho, L.T.Y., Wei, Y., Fogarty, N.M.E., Blakeley, P., Sherwood, R.I., Ji, H., and Niakan, K.K. (2015). Gata6 potently initiates reprograming of pluripotent and differentiated cells to extraembryonic endoderm stem cells. Genes Dev. *29*, 1239–1255. https://doi.org/10.1101/gad.257071.114.

Zhang, Y., Park, C., Bennett, C., Thornton, M., and Kim, D. (2021). Rapid and accurate alignment of nucleotide conversion sequencing reads with HISAT-3N. Genome Research *31*, 1290–1295. https://doi.org/10.1101/gr.275193.120.
